# SARS-CoV-2 Omicron Spike Glycoprotein Receptor Binding Domain Exhibits Super-Binder Ability with ACE2 but not Convalescent Monoclonal Antibody

**DOI:** 10.1101/2021.12.09.471885

**Authors:** I. O. Omotuyi, O. Olubiyi, O. Nash, E.O. Afolabi, B.E. Oyinloye, S. Fatumo, M. N. Femi-Oyewo, S.E. Bogoro

**Affiliations:** Institute for Drug Research and Development, S.E. Bogoro Center, Afe Babalola University, Ado Ekiti, Nigeria; Department of Pharmaceutical Chemistry, College of Pharmacy, Afe Babalola University, Ado-Ekiti, Nigeria; Molecular Biology and Molecular Simulation Center (Mols&Sims), Ado Ekiti, Nigeria; Centre for Genomics Research and Innovation, National Biotechnology Agency, Nigeria; London School of Hygiene and Tropical Medicine, London, United Kingdom

## Abstract

**Background:** SARS-CoV-2, the causative virus for COVID-19 has now super-mutated into the Omicron (*Om*) variant. On its spike glycoprotein alone, more than 30 substitutions have been characterized with 15 within the receptor binding domain (RBD); It therefore calls to question the transmissibility and antibody escapability of Omicron. This study was setup to investigate the Omicron RBD’s interaction with ACE2 (host receptor) and a SARS-CoV-2 neutralizing monoclonal antibody (mAb).

**Methods:** *In-silico* mutagenesis was used to generate the Om-RBD in complex with ACE2 or mAb from the wildtype. All-atom molecular dynamics (MD) simulation trajectories were analyzed for interaction.

**Results:** MD trajectories showed that Omicron RBD has evolved into an efficient ACE2 binder, via *pi-pi* (*Om*-RBD-Y501/ACE2-Y41) and salt-bridge (*Om*-RBD-K493/ACE2-Y41) interactions. Conversely, in binding mAb, it has become less efficient (Center of mass distance of RBD from mAb complex, wildtype ≈ 30 Å, Omicron ≈ 41 Å). Disruption of *Om*-RBD/mAb complex resulted from loose interaction between *Om*-RBD and the light chain complementarity-determining region residues.

**Conclusions:** Omicron is expected to be better transmissible and less efficiently interacting with neutralizing convalescent mAbs.

**General significance:** Our results elucidate the mechanisms for higher transmissibility in Omicron variant.

## Main Body

COVID-19 (coronavirus disease 2019) is caused by the novel coronavirus severe acute respiratory syndrome-coronavirus-2 (SARS-CoV-2) (Zhou et al. 2020). SARS-CoV-2 tropism is initiated when its spike (S) glycoprotein binds to the host angiotensin-converting enzyme 2 (ACE2) and its partner transmembrane serine protease 2 (TMPRSS2) (32142651) serving as door-way to cellular entry. In addition to its role as the receptor (Wrapp et al., 2020), the S protein is also the key targets for several antibodies currently in use as treatment options for COVID-19; especially the receptor binding domain (RBD) (Zhou et al., 2020).

Notably, the antibodies generated by the Pfizer-BioNTech mRNA-BNT162b2, Moderna mRNA-1273 ultimately targets SARS-CoV2 spike glycoprotein (**Fig. *1a, b-ii*)** at the receptor-binding domain (RBD, **Fig. *1b, i-ii***)) while Astra-Zeneca-ChAdOx1-S and Janssen-Ad26.COV2.S were primarily designed to express SARS-CoV-2 spike protein as immunogen (Mascellino et al., 2021). Curiously, many convalescent monoclonal antibodies (mAbs) bind at the ACE2 site on the RBD (Bertoglio et al., 2021). It is therefore not surprising that as SARS-CoV-2 variants with mutations in the RBD begin to emerge (Omotuyi et al., 2020), so is concern over transmissibility and antibody escape (Mascellino et al., 2021). Variants whose RBD mutations eventually resulted in worse clinical outcomes include: B.1.1.7 (N501Y), B.1.351/P1 (K417T/N, E484K, N501Y) (Queiros-Reis et al., 2021), and B.1.617.2 (Delta variant; L452R, T478K); then the most recent (B.1.1.529), also termed the Omicron variant. (Gao et al., 2021). Intriguingly most S protein substitutions (N440K, G446S, S477N, T478K, E484A, Q493R, G496S, Q498R, N501Y, Y505H, **Fig. *1c***) occur at the ACE2-binding site of the RBD (**Fig. *1d***).

**Figure 1.0.**
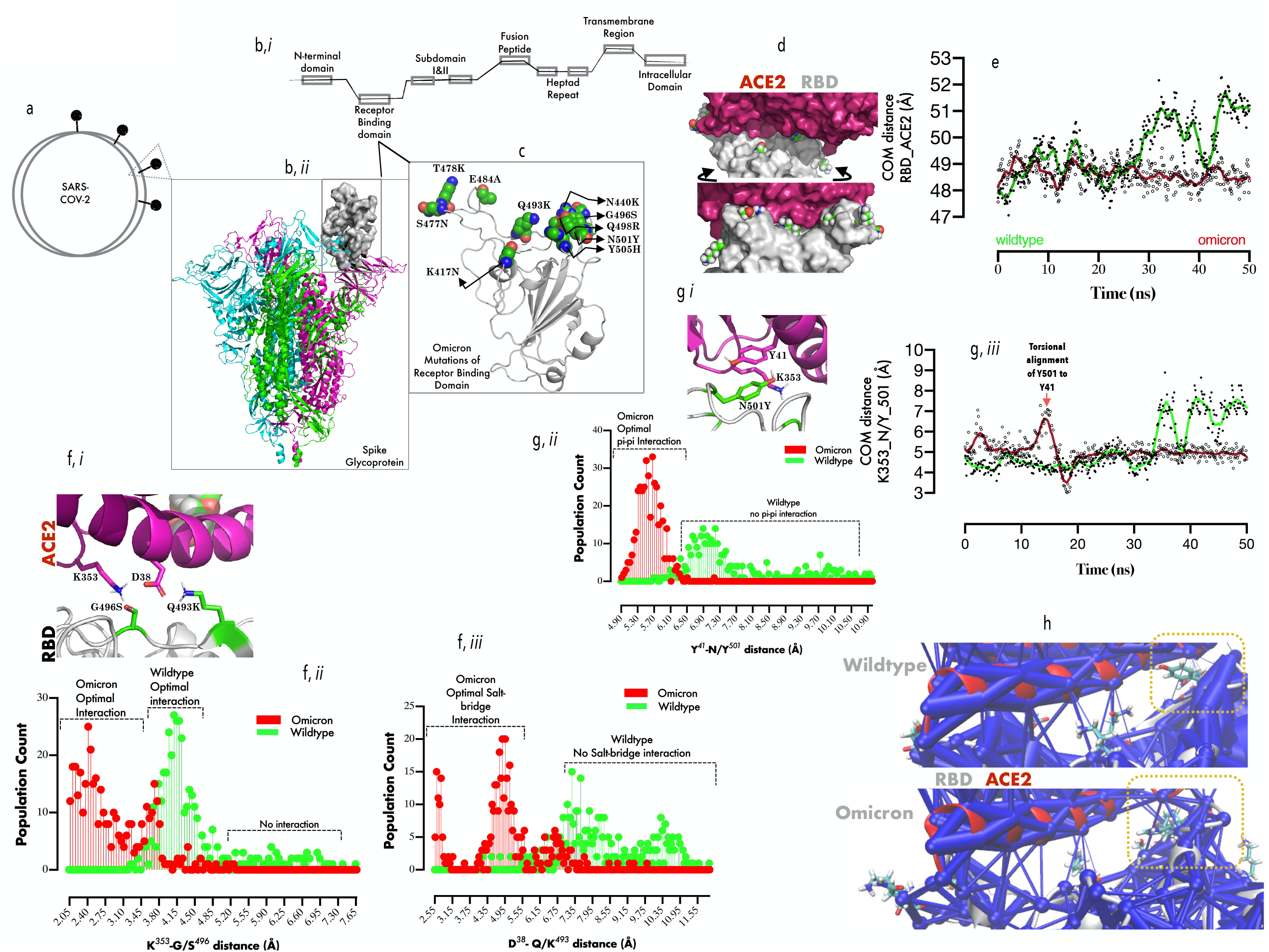
Comparative binding dynamics of Omicron and Wildtype RBDs to ACE2: 1a: Representation of SARS-CoV-2 and S protein, (b, i) Representation of the different constituent regions in a typical monomeric S protein. (b, ii) Cartoon representation of the trimeric S protein, showing one of the three RBDs in up configuration (surface representation). (c). Cartoon representation of the RBD, showing the cluster of substitutions (represented in vmd spheres) that define the Omicron variant. (d, upper and lower plane) Surface representation of ACE2/RBD complex showing the spatial distribution of the Omicron substitutions around the RBD. (e) Smoothened line graph showing the mean center of mass distance between RBD and ACE3 with time. (f, i-iii) Spatial projection of ACE2-K353/D38 proximal to the RBD-G966S/Q493K (i), and population count distributions of side-chain atom distance between K353/G966S (ii) and D38/Q493K (iii). (g, i-iii) Spatial projection of ACE2-K353/Y41 proximal to the RBD-N501Y (i), and population count distributions of side-chain atom distance between Y41/Y501 (ii) and time-evolved smoothened mean distance between K353 and Y501 (iii). (h) The network data showing weighted interaction between ACE2/RBD in wildtype (upper plane) and Omicron (lower plane). Yellow rectangles indicate the substitution cluster and their effect on the weight of RBD-ACE2 interaction.

First, in order to provide insight into how the substitutions affect ACE2 binding, all-atom MD simulation in explicit water was set up (Supplementary ***Methods***) and checked for important biological events every 10 ns. Surprisingly, Wildtype-but not Omicron RBD exhibit intermittent dissociation from ACE2. The largest amplitude of dissociation occurs at 30 ns, (*Om-*vs *Wt-*: ≈ 48.5 Å *vs* 51.5 Å) and 45 ns (*Om-*vs *Wt-*: ≈ 48.0 Å *vs* 52.5 Å, **Fig. *1e***), indicating that Omicron, but not the wildtype RBD has better ACE2 binding capacity.

Whilst it is to be noted that previous reports identified T478K, and N501Y substitutions are associated with increased ACE2 binding (Kim et al., 2021), in Omicron RBD, S496 resides within hydrogen-bond distance (≈ 3.5 Å) from ε-amino group of ACE2-K353 (**Fig. *1f, i***), which is absent in Wildtype (G496, distance > 4.5 Å, **Fig. *1f, ii***). Omicron K493 also evolved salt-bridge interaction with ACE2 D38 (distance ≈ 2.5-4.5 Å) as opposed to Q493 which fails to form hydrogen bond (distance > 6.0 Å, **Fig. *1f, ii***). Further investigation showed that G496 (wildtype) allowed ACE2-K353/D38 salt-bridge interaction rather than engaging RBD (data not shown), thus, further weakening ACE2 binding in the wildtype.

We further elucidate that the stabilizing effect of N501Y mutation on ACE2 occurs through ACE2-K353 interaction (**Fig. *1g, i***). Here, the measurement of the inter-atomic distance between the phenolic side chains of Y501 (RBD) and Y41 (ACE2) indicated a possible *pi-pi* interaction (distance < 6.0 Å, **Fig. *1g, ii***); a feature non-existent in wildtype (N501, distance > 6.9 Å, **Fig. *1g, ii***). Y501/41 stacking spatially locks Y501 in place allowing cation-pi interaction (**Fig. *1g, ii***) with K353 (ACE2). A representation of the difference in the strengths of ACE2 interaction offered by wildtype (**Fig. *1h*, *upper plane***) and Omicron (**Fig. *1h*, *lower plane***) RBDs were also projected using a weighted network representation, with Omicron RBD residues displaying stronger network interaction within the substitution clusters. Without a doubt, Omicron S protein RBD exhibits super-binder ability with ACE2 with resulting higher transmissibility potentials.

Next, we sought to understand how the Omicron RBD substitutions affect mAb binding, protein-protein docking and scoring with several available antibodies was initially performed. A cursory look suggests that Omicron binding is associated with improved binding scores (Supplementary **Table 1.0)** but when one of the complex (Bertoglio et al., 2021) was subjected to simulation, the events were different. First, the mean COM distance separating RBD from the mAb complex (heavy & light chains, **Fig. 2a, *i***) over the entire trajectories showed that the Omicron was more loosely bound to the mAb in comparison with wildtype (*Om-*vs *Wt-*: ≈ 41.0 Å *vs* 30.0 Å, **Fig. 2a, *ii***), and we showed that the antigen-binding fragments (Fab, residues 1-108 (heavy chain), residues 1-106 (light chain), **Fig. 2a, *iii***), for the first 30 ns was ≈ 3 Å less compact in binding Omicron RDB in comparison with the wildtype (**Fig. 2a, *iv***), This result suggest that local events at the RBD-Fab interface is responsible for the difference in mAb binding. Therefore, we further investigated the roles of each chain as most of the mutations cluster at the VLCDR binding interface (**Fig. 2b, *i***). A population plot of the COM distance between the mAb-light chain and RBD strongly suggest that Omicron RBD binds beyond sub-optimal distance (distance > 50 Å, **Fig. 2b, *ii***) in comparison with the wildtype (distance < 47 Å) and surprisingly a similar pattern was observed in the heavy chain (Wildtype ≈ 37.5 Å *vs*. Omicron ≈ 38.5 Å, **Fig. 2b, *iii***).

**Figure 2.0.**
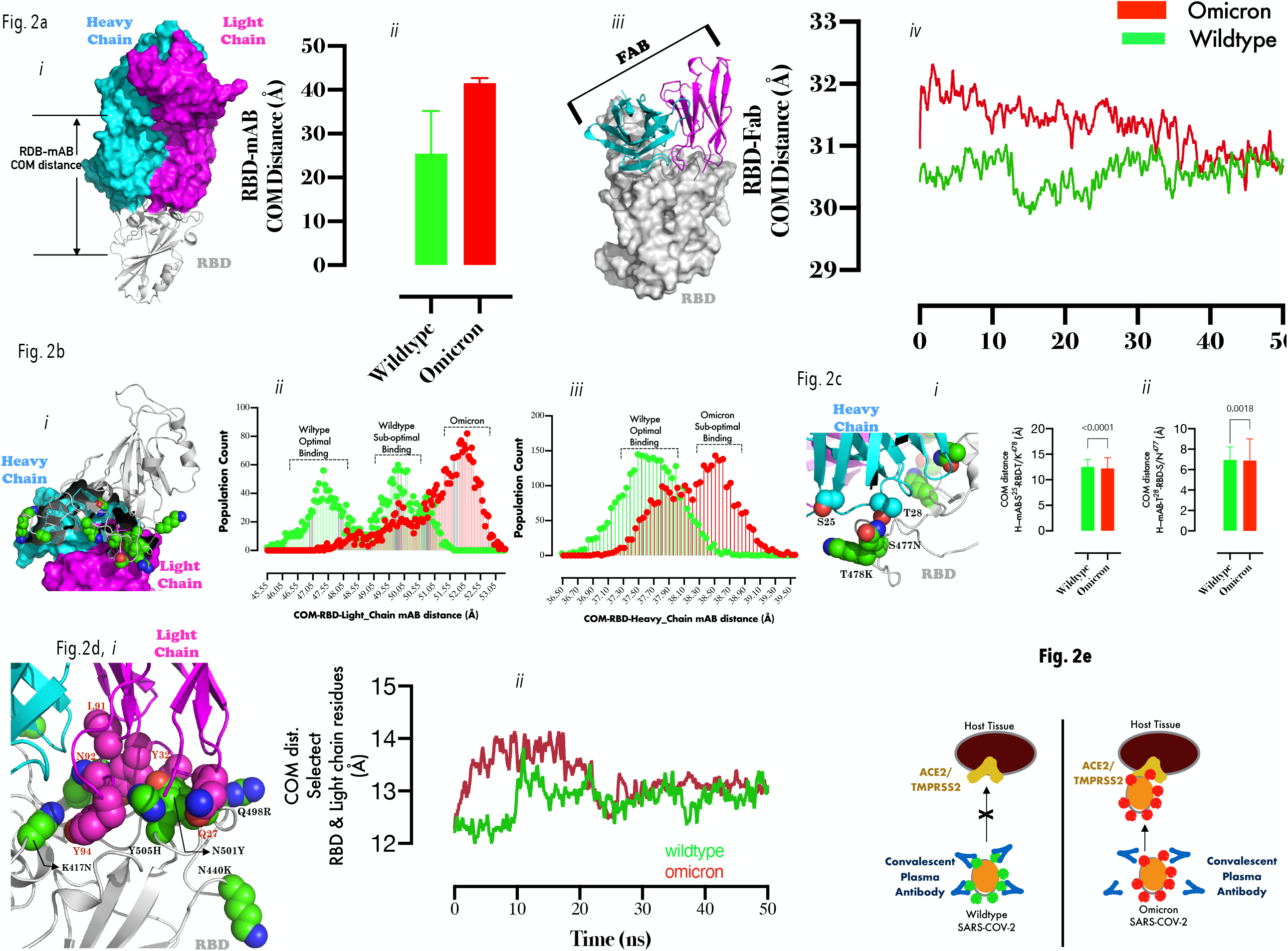
Comparative binding dynamics of Omicron and Wildtype RBDs to convalescent mAb: 2a (i) Representation of convalescent mAb (blue surface=heavy chain, pink surface=light chain) in RBD bound (gray cartoon) state. (a, ii) The bar chart plot of the COM distance between RBD and mAb during MD simulation. (a, iii) Representation of MAb-Fab (cartoon)/RBD(surface) complex. (2, iv) Smoothened line graph of showing the mean center of mass distance between RBD and Fab with time. (2b, i) A spatial representation of Omicron RBD substitutions relative to the light and heavy chains of mAb. (2b,ii) population count distributions of COM distance between the light chain/RBD and heavy-chain/RBD (iii). (2c, i) Zoomed representation of heavy chain CDR loop residues (S25/T28) proximal to some substituted RBD residues (T478K, S477N). (2c, ii) Bar graph plots of the inter-residue distance between S25 and T/K478(ii) and T28/S/N477 (T-test comparison were made at p< 0.05 using two-tailed paired sampling and parametric test). (2d, i) Zoomed representation of light chain CDR loop residues (Q27, Y32, L91, N92, and Y94) proximal to substituted RBD residue cluster (K417N, N440K, Q498R, Y505H). Smoothened line graph showing the mean center of mass distance between the clustered RBD substitutions and the light chain CDR loop residues with time. Fig. 2e: A mechanistic projection showing a tightly bound convalescent mAbs to Wildtype SARS-Cov-2-RBD thus, protecting ACE2/TMPRSS2-experessing cells from infection; while loosely bound convalescent mAbs to Omicron SARS-Cov-2-RB coupled with high-potency interaction between ACE2 and Omicron-RBD promotes higher infectivity in ACE2/TMPRSS2-experessing cells.

Finally, specific mAb residues which account for the sub-optimal binding were identified. Loss of interaction between S25/T28 (heavy chain CDRs) and T478K and S477N respectively (**Fig. 2c, *i*, *right and left panels***) partially explain the loose binding with Omicron RBD and time-evolved dynamics setup for to monitor the interaction between clustered RBD substitutions (K417N, N440K, Q493R, G496S, Q498R, N501Y, and Y505H) and the VLCDR loop residues (**Fig. 2d, *i***,). The 2 Å separation of VLCDR loop residues from Omicron RBD for the first 20 ns of simulation (reconverged afterwards) is consistent with a previous studies where N501Y and K417N were associated with detached RBD from mAb light chain CDR1 loop (Dejnirattisai et al., 2021). These results suggest strongly that Omicron RBD is ACE2 super binder but damped convalescent mAb binder (**Fig. 2e**).

## Notes

### Competing Interest Statement

The authors have declared no competing interest.

