## Supplementary_Methods_Figure for "SARS-CoV-2 Omicron Spike Glycoprotein Receptor Binding Domain Exhibits Super-Binder Ability with ACE2 but not Convalescent Monoclonal Antibody"

Supplementary Table 1.0:

**Docking Scores (Kj/mol) of Omicron and Wildtype RBDs in mAB bound Complexes**

| PDB IDs | Antibody Designation | Wildtype | Omicron |
| --- | --- | --- | --- |
| 7DEO | PR1077 | -424.48 | -455.43 |
| 7DEU | PRO953 | -239.41 | -247.09 |
| 7DET | PRO961 | -317.08 | -341.44 |
| 7CJF | P4A1 | -215.27 | -243.05 |
| 7B3O | STE90-C11 | -305.29 | -398.53 |

**Starting Structures:**

**RBD-ACE2 Complex:**

Wildtype RBD in ACE2 bound state previously resolved (PDB ID: 7KMB) (Zhou et al. , 2020) was retrieved. All broken chains and incomplete residues were reconstructed in protein preparation module of Schrodinger Suite. In order to generate the Omicron RBD, the following substitutions (N440K, G446S, S477N, T478K, E484A, Q493R, G496S, Q498R, N501Y, Y505H) were made using PyMol Mutagenesis plugin.

**RBD-mAB Complex:**

Different neutralizing mABs bound to unique regions of the RBD previously deposited: 7DEO/7DEU/7DET (Fu et al. , 2021), 7CJF (Guo et al. , 2021), 7B3O (Bertoglio et al. , 2021) were retrieved. Prior to antibody re-docking experiment, all broken chains and incomplete residues were reconstructed in protein preparation module of Schrodinger Suite. Omicron RBD was generated using PyMol Mutagenesis plugin.

**Antibody Docking**

The HDOCK server (Yan et al. , 2020) for integrated protein–protein docking was used to reproduce crystallographic poses and scoring of the RBD-mAB poses for both wildtype and Omicron.

**Biosystem generation for Atomistic Simulation**

To generate RBD-ACE2 (PDB ID: 7KMB) or RBD-mAB (PDB ID: 7B3O) biosystems for Omicron and Wildtype for simulation, CHARMM-GUI webserver ([www.charmm-gui.org](http://www.charmm-gui.org)) (Jo et al. , 2008) was used. All protein were parameterized in CHARMM36 all-atom additive protein force field (Huang and MacKerell, 2013) while glycan parameterization was performed using ParamChem service (https://cgenff.paramchem.org) as implemented on CHARMM-GUI webserver interface. Each biosystem was solvated in TIP3P explicit water model (Florova et al. , 2010) and neutralized with Na^+^/CL^−^ .

**Molecular dynamics (MD) simulation**

All molecular dynamics simulation was run on NAMD molecular dynamics software (Phillips et al. , 2005) in three stages of minimization, equilibration and production. During equilibration, the biosystems were under constant pressure and temperature (NPT; 298K, 1 bar) conditions using Berendsen temperature and pressure coupling algorithms. All Van der Waals interactions were estimated at 10 Å, while electrostatic interactions were estimated using particle mesh Ewald (PME) summation equation and equation of atomic motion was integrated using the leap-frog algorithm at 2 fs time step for a total time of 30 ns with positional restraints imposed on the heavy atoms in all directions.

In order to generate the two independent states for production stage MD simulation, equilibration stage trajectories were loaded into VMD (Humphrey et al. , 1996) and two structures with the largest rmsd were retrieved and simulated as discussed for equilibration above for 50 ns with the removal of restraints. All trajectories were checked for convergence simulations, lipid bilayer thickness was fairly maintained between 3 ~ 4 nm throughout the (data not shown) prior to analyses. All calculations were performed on SuperMicro workstations (32-E2600 Intel Xeon CPUs, 2 M 6000 GPUs Accelerator PCI-E x16 Card/node) housed at the S.E. Bogoro Center, Afe Babalola University, Ado-Ekiti, Nigeria.

**Post-MD simulation analyses and data presentation**

Dynamical networks for RBD-ACE2 interaction for both wildtype and Omicron systems were calculated as described (Sethi et al. , 2009), we have previously described the use of *Carma* (ver. 1.4), *gncommunities* and *subopt* scripts for generating files for network analysis (Omotuyi et al. , 2015). Network tools implemented in VMD was used to visualize the source-sink pairs. A pair of nodes was connected by an edge if the corresponding residues were resident within 4.5 Å distance for at least 80% of the frames analyzed while the edge size is weighted.

Unless otherwise stated, all inter-group, inter-residue or inter-atomic distances were calculated using PLUMED plugin for molecular dynamics (Bussi and Tribello, 2019).

All line graphs, bar charts, or population counts were plotted as mean from 2  independent runs using GraphPad prism (ver 9.0).
